## Supporting Information for "Cortical feedback and gating in odor discrimination and generalization"

### Pattern convergence and divergence for normal distributions of module responses: numerical results

We noted that the firing rate of modules should be determined by the sum of the firing rates of the component mitral cells. Thus, by the central limit theorem, the module firing rates should be normally distributed. Likewise, feedback affects individual mitral cells, and the resulting change in the firing rate of a module is determined by the sum of these changes for the component mitral cells. Thus, the effect of feedback on the module firing rates should also be normally distributed. We then simulated $N = 10,000$ cortical units receiving inputs from the corresponding number of bulb modules, and responding to two odors A and B. Before feedback, we took the responses of bulb modules to be normally distributed for each odor. We controlled initial odor similarity by changing the fraction of modules that responded to both odors as opposed to only one. We then modeled the effects of contextual feedback by adding normally distributed firing rate changes to the modules. The mean and standard deviation of the distributions were picked so that the range of module responses would be comparable to the previous analytical computations. Additionally, the hard threshold for cortical activation in the analytical model was replaced with a sigmoid acting on the olfactory bulb module responses with a soft threshold at $\theta_c$. These simulations of the statistical model produced trends in pattern convergence and divergence that were qualitatively similar to those derived analytically. We also were able to show that these results were robust to broad changes in parameters (Fig. S1).

### The necessity of a nonlinear transfer function with high cortical threshold in simulations of the statistical model

Our results depend critically on a two-layer architecture where activity in the bulb is modulated by feedback and then passed through a nonlinearity with a high threshold in the cortex. To see this, we changed the nonlinearity to a linear function $f(R_i) = R_i$ and

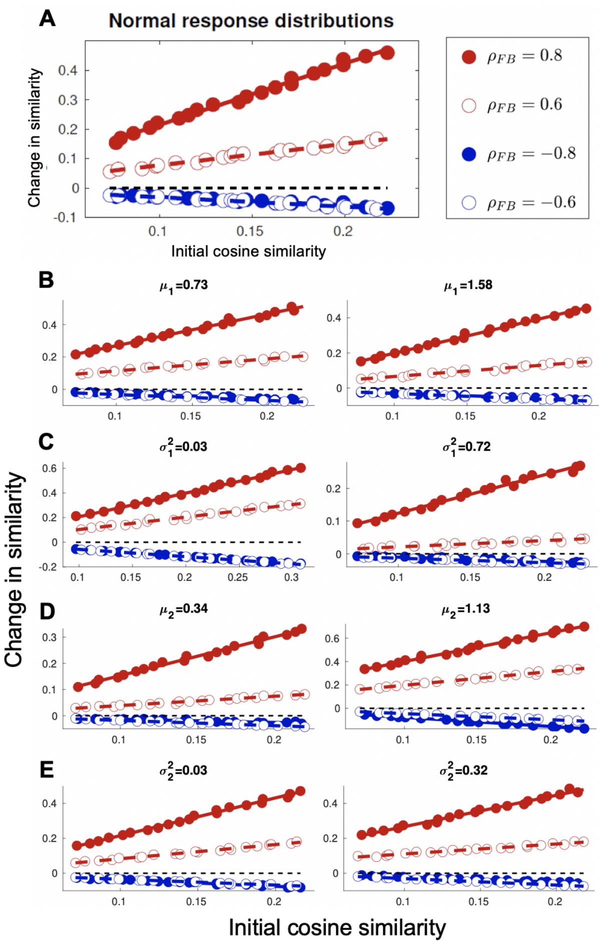


**Figure S1.** The trends in pattern convergence and divergence are robust against changes in the parameters of the normal distributions of module responses. (**A**) Results of the analytical model generalize to realistic normal distributions of module responses to odor and feedback inputs (mean and variance $\mu_1 = 1.15$, $\sigma_1^2 = 0.42$ for odor inputs; and $\mu_2 = 0.57$, $\sigma_2^2 = 0.28$ for feedback inputs). The cortical threshold is set to $\theta_c = 1.6$ to give cortical activation of 10% [1]. Datapoint obtained by averaging the results for 10 randomly generated pairs of odor inputs with similar similarity values. Different markers indicate different feedback conditions. Filled red/blue circles: the two feedback scenarios of Fig. 1 in the main text, corresponding to $\rho_{FB} = 0.8$ and $\rho_{FB} = -0.8$, respectively. Note that $|\rho_{FB} \neq 1|$ due to variability in the amplitude of the feedback strength $\Delta R$. Empty red/blue circles: two intermediate conditions, corresponding to 50% of modules receiving feedback, of which 75% are shared between the two odors. Of the feedback-targeted modules, 75% receive inhibitory feedback for both odors in the first case (red, $\rho_{FB} = 0.6$), and 75% / 12.5% receive inhibitory feedback for the first/second odor in the second case (blue, $\rho_{FB} = -0.6$) (**B–E**) Each panel shows the statistical model results assuming normal distributions of module responses to odor and feedback inputs. The parameters of the distributions are varied one by one with respect to **A** ($\mu_1 = 1.15$, $\sigma_1^2 = 0.42$, $\mu_2 = 0.57$, $\sigma_2^2 = 0.28$). Results are robust against changes in (**B**, **C**) the mean and variance, respectively, of the distributions of module responses to odor inputs and (**D**, **E**) the mean and variance, respectively, of the distributions of module responses to feedback inputs. As in **A**, each datapoint is obtained by averaging the results for 10 randomly generated pairs of odor inputs with similar similarity values. Different markers indicate different feedback conditions. Filled red/blue circles: the two feedback scenarios of Fig. 1 in the main text, corresponding to, respectively, $\rho_{FB} = 0.8$ and $\rho_{FB} = -0.8$ in **B–C**, $\rho_{FB} = 0.75$ and $\rho_{FB} = -0.75$ in **D, left**, $\rho_{FB} = 0.94$ and $\rho_{FB} = -0.94$ in **D, right**, $\rho_{FB} = 0.92$ and $\rho_{FB} = -0.92$ in **E, left**, $\rho_{FB} = 0.72$ and $\rho_{FB} = -0.72$ in **E, right**. Empty red/blue circles: two intermediate conditions, corresponding to 50% of modules receiving feedback, of which 75% are shared between the two odors. Of the feedback-targeted modules, 75% receive inhibitory feedback for both odors in the first case (red, $\rho_{FB} = 0.6$ in **B–C**, $\rho_{FB} = 0.56$ in **D, left**, $\rho_{FB} = 0.71$ in **D, right**, $\rho_{FB} = 0.69$ in **E, left**, $\rho_{FB} = 0.54$ in **E, right**), and 75% / 12.5% receive inhibitory feedback for the first/second odor in the second case (blue, $\rho_{FB} = -0.6$ in **B–C**, $\rho_{FB} = -0.56$ in **D, left**, $\rho_{FB} = -0.71$ in **D, right**, $\rho_{FB} = -0.69$ in **E, left**, $\rho_{FB} = -0.54$ in **E, right**)

studied how the similarity in bulb responses to odors changes due to feedback. As above, we assumed that these responses prior to feedback and the feedback-induced changes were both normally distributed, and we also maintained the same feedback similarities and statistics. We then computed the cosine similarity between bulb responses to odors A and B, before and after adding feedback. Fig. S2A shows that the effects induced by feedback in the bulb are substantially different from those that arise in the cortical layer. In particular, pattern convergence can be achieved only when the feedback similarity is very high, and even then decreases with increasing odor similarity, oppositely to the trend in the cortical layer. Moreover, the effects induced by feedback in a two-layer network with a low cortical threshold (and normally distributed responses) are again qualitatively different from those arising at high cortical thresholds and resemble closely the results for low cortical thresholds found analytically in Fig. 4A (Fig. S2B). Thus, the form of pattern convergence of Fig. S2C can only emerge in a network with layers linked by a non-linear, high-threshold transfer function.

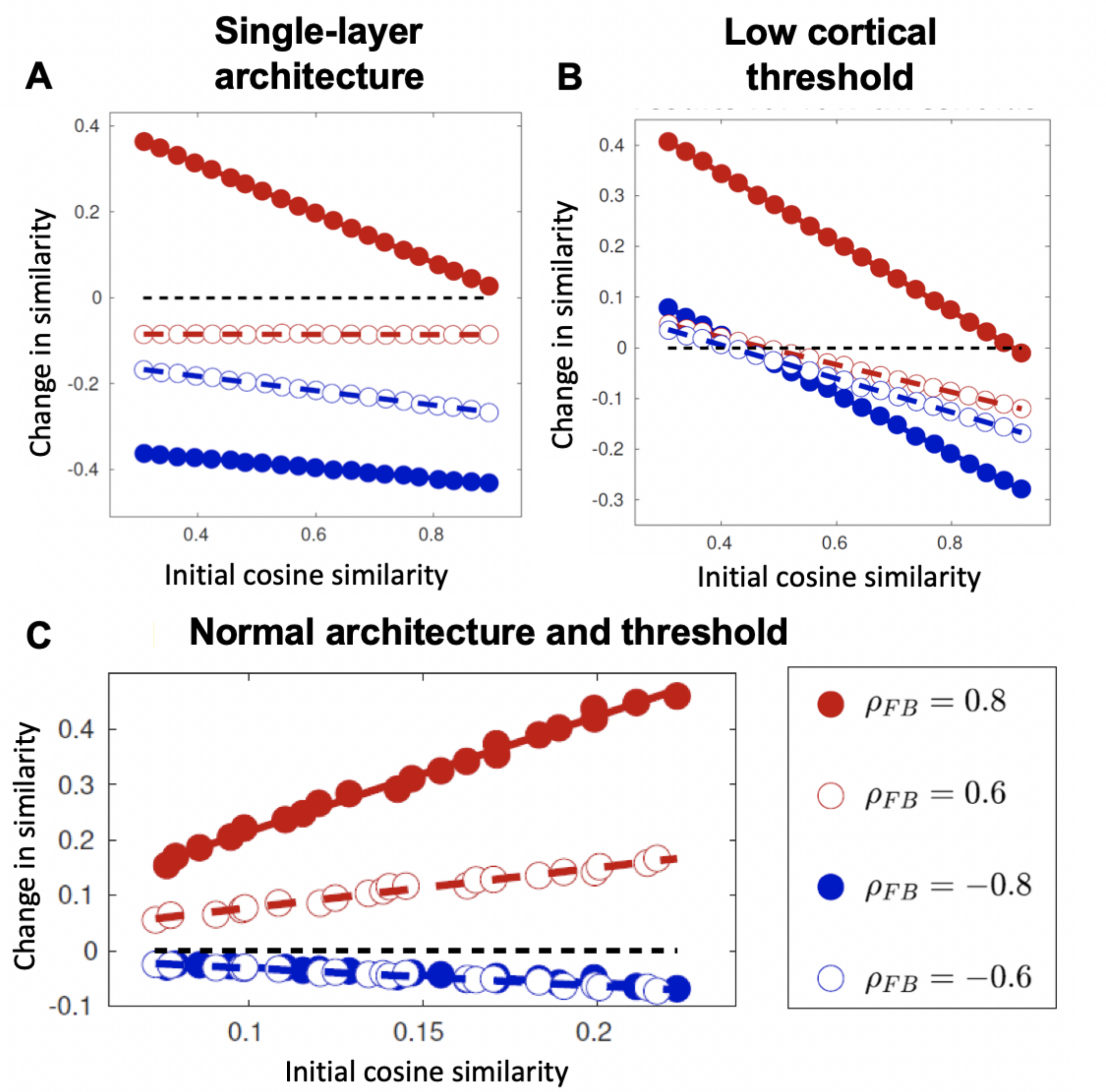


**Figure S2.** The mechanism for pattern completion and separation requires a network with (at least) two layers linked by a high-threshold transfer function. The statistical effects induced by feedback in a single-layer architecture or with low cortical threshold are qualitatively different from those arising in a two-layer model with a high-threshold transfer function. (**A**) With only one layer, pattern completion can be achieved only when the feedback similarity is very high and decreases with increasing odor similarity (red). Moderately correlated and anticorrelated feedback induce similar effects (empty red and blue circles). Same conditions as in **C**: Filled red/blue circles for the two extreme feedback scenarios with $\rho_{FB} = 0.8$ and $\rho_{FB} = -0.8$, respectively; empty red/blue circles for the two intermediate feedback conditions, with $\rho_{FB} = 0.6$ and $\rho_{FB} = -0.6$, respectively. (**B**) Same conditions as in **C** except with lower cortical threshold. The trend reversal is similar to that seen in the analytical framework (Fig. 4A). (**C**) Figure S1A demonstrating results from a normal two-layer, high-threshold architecture for comparison.

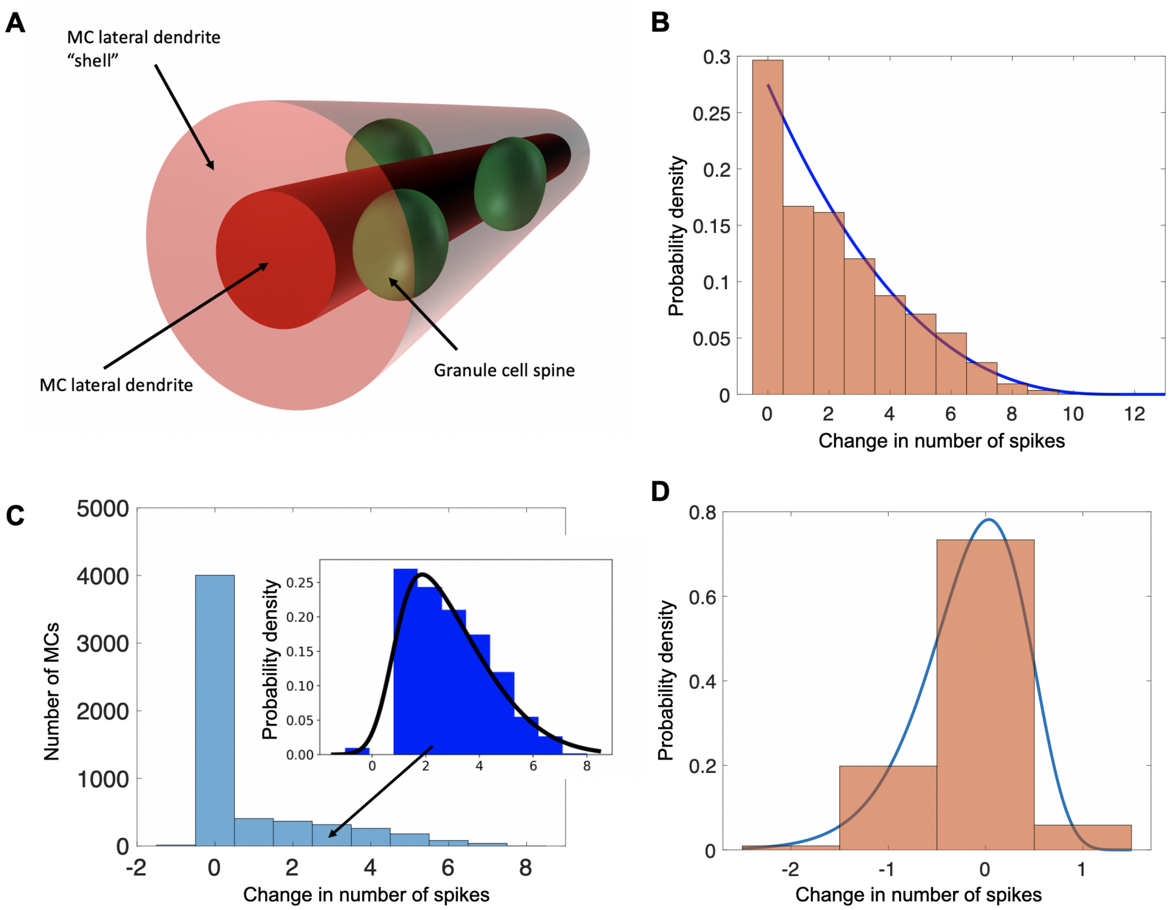


**Figure S3.** Details of the biophysical model and subsequent distributions in firing rates. (**A**) Depiction of the relevant volumes for calculating MC-GC connectivity (**B**) Distribution of MC firing rates due to odor input (**C**) Distribution of changes in MC firing rates due to excitatory feedback. Inset shows the skew normal distribution that was sampled if the change was not equal to 0. (see "Firing rate distributions") (**D**) Distribution of changes in odor-receiving MC firing rates due to excitatory feedback to GCs. Note that the lognormal distribution has been shifted and flipped to match the range of the data. The parameters of the simulations for the distributions in **B**–**D** are provided in the text of the Methods ("Neuronal and network dynamics; Firing rate distributions")

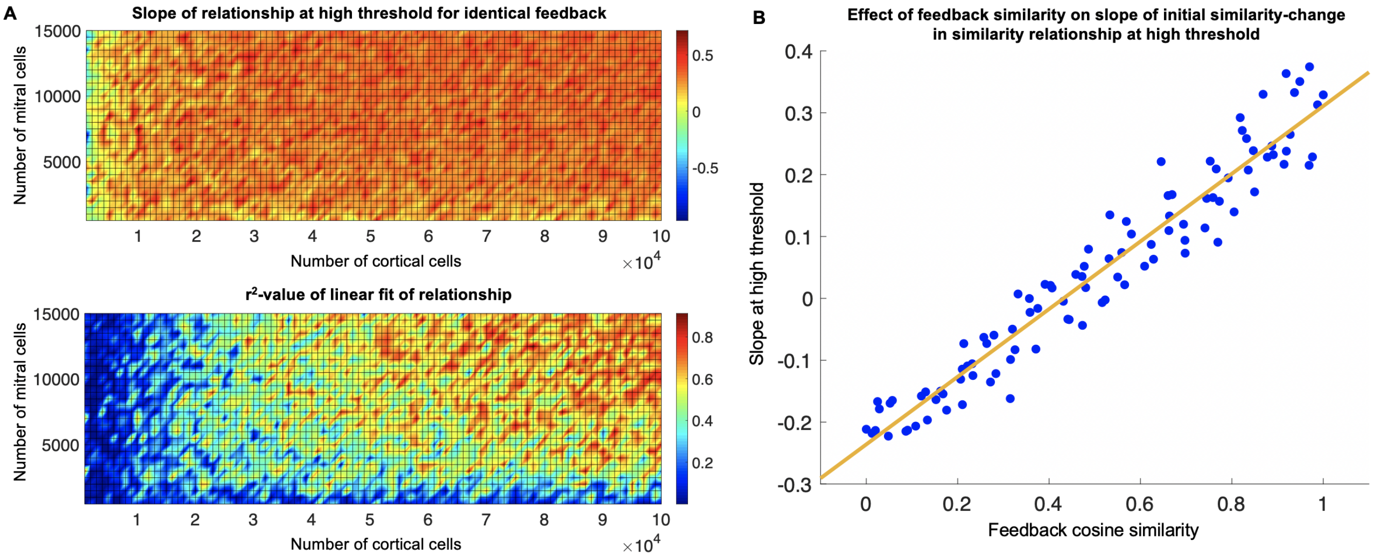


**Figure S4.** Relationship between initial similarity and change in similarity at high threshold in the mechanistic model. (**A**) A robustly positive slope of the relationship between initial similarity and change in similarity at high threshold is only achieved for a sufficient number of MCs and bulb modules (i.e., cortical cells). For different numbers of $M$ mitral cells and $C$ bulb modules, we simulated presentation of the same positive feedback for different pairs of odors and then measured the slope of the relationship between initial similarity and change in similarity at high threshold. Although positive slope was achieved for relatively low numbers of MCs and bulb modules, this relationship did not achieve a consistently high $r^2$ value without approximate values of $M > 8000$ and $C > 80000$. (**B**) Sufficiently correlated positive feedback produces a proportional relationship between initial similarity and change in similarity. We generated a range of feedback similarities for pairs of excitatory feedback vectors. We found that the slope of the relationship between initial similarity and change in similarity at high threshold varied linearly with the feedback similarity (slope $= 0.5469$, $r^2 = 0.9322$). Thus, for feedback vectors with significant similarity, the relationship between initial similarity of two odor representations and the change in similarity of those representations following feedback is positive. For all simulations, $M = 10000, K = 100000, f_{odor} = 0.12, G = 500, p_{FB} = 0.08$, and $q = 0.07$.
